# Motor-driven modulation of actin network mechanics across linear and nonlinear regimes

**DOI:** 10.1101/2025.06.15.659806

**Authors:** Bernard Nkusi, Morgan Thomas, Bekele J. Gurmessa

## Abstract

Cytoskeletal networks enable cells to dynamically regulate their mechanical properties in response to internal forces and external cues. Here, we investigate how motor activity influences the structure and mechanics of actomyosin networks reconstituted *in vitro* from filamentous actin, myosin II minifilaments, and transient *α*-actinin cross-linkers. By varying the myosin-to-actin molar ratio (*R*_MA_), we observe a transition from isotropic actin meshes to contractile, coarsened architectures marked by bundled filaments and increasing spatial correlation lengths (*ξ*_z_, *ξ*_t_). Optical tweezers microrheology reveals a nonmonotonic mechanical response: at low *R*_MA_, networks fluidize, with reductions in the plateau modulus (*G*^0^), zero-shear viscosity (*η*^0^), and fast relaxation timescales (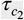, *τ*_1_). At higher motor levels, the networks stiffen and retain internal stress, reflecting contractile reinforcement. Notably, *τ*_1_ exhibits a minimum when plotted against *ξ*_z_, suggesting that intermediate levels of coarsening facilitate efficient local stress dissipation. These results identify distinct mechanical regimes governed by motor-induced remodeling and highlight a structural basis for the dual roles of myosin in fluidization and reinforcement.

## 1 INTRODUCTION

The eukaryotic cytoskeleton is a dynamic, multifunctional scaffold composed of actin filaments, intermediate filaments, and microtubules that collectively maintain cellular architecture, enable shape change, and regulate internal organization [1, 2]. This system supports a diverse array of vital cellular processes, including migration, division, morphogenesis, and mechanotransduction [3, 4, 5, 6, 7, 8, 9, 10].

Central to these mechanical roles are networks of filamentous actin (F-actin), which assemble through ATP-dependent polymerization of globular actin (G-actin) monomers into semiflexible filaments [3]. These filaments self-organize into complex architectures–branched, bundled, or entangled–that support protrusive structures like filopodia and lamellipodia, crucial for cellular motility and signaling [11, 12, 13]. The mechanics and organization of actin networks are finely tuned by actin-binding proteins (ABPs), including passive crosslinkers such as *α*-actinin and active force generators like myosin II [14, 15], as well as by electrostatic interactions and counterion effects [16, 17, 18, 19].

Crosslinking proteins, whether transient or permanent, fundamentally shape the mechanical and rheological properties of actin networks by regulating filament connectivity, bundling, and stress dissipation pathways. Transient crosslinkers such as *α*-actinin, filamin A, fascin, and myosin II (under physiological ATP conditions) reversibly bind actin filaments with lifetimes on the order of seconds, enabling networks to stiffen moderately under small strains while still permitting stress relaxation and dynamic remodeling [11, 20, 21, 22, 23]. *α*-Actinin, for instance, plays a pivotal role in cytokinesis by regulating contractile ring dynamics [24] and shapes inter-phase cell mechanics by promoting stress fiber formation and mediating strain stiffening [11, 25, 24, 26]. In contrast, permanent crosslinkers such as biotin–NeutrAvidin crosslinker form effectively a very strong bond between filaments, dramatically enhancing the elastic modulus while suppressing dissipative modes, producing networks that resist stress relaxation and exhibit network rupturing at high strains [27, 28, 29, 30].

Myosin II, one of the most studied actin binding proteins [31, 32, 33, 34, 35, 36, 37], provides a unique dual role: it not only transiently crosslinks actin but also actively generates contractile stresses via ATP-driven filament sliding, assembling into minifilaments that orchestrate large-scale network reorganization far from equilibrium [38, 39, 40, 21, 41, 15, 42]. Notably, under low ATP conditions, myosin II can shift into a passive crosslinking role, enhancing filament connectivity and mechanical resilience, as revealed by microrheology measurements showing increased plateau moduli (*G*^0^) and altered stress relaxation behavior [34, 36, 21, 43].

These crosslinkers critically modulate microrheological properties, including the elastic modulus *G*^′^(*ω*), the viscous modulus *G*^′′^(*ω*), and the characteristic relaxation timescale *τ*_*c*_, as well as the nonlinear mechanical response under strain. Transient crosslinkers typically increase *G*^′^ but allow *G*^′′^ to dominate at low frequencies, reflecting viscoelastic fluidity, while permanent crosslinkers push the system toward an elastic-dominated response by elevating *G*^′^ and suppressing *G*^′′^ across the frequency spectrum [44, 45, 27, 46]. Relaxation times *τ*_*c*_ become longer in permanently crosslinked systems due to restricted filament mobility, whereas transiently crosslinked or motor-active networks often display faster relaxation pathways owing to ongoing remodeling [21, 15, 47, 48]. Nonlinear microrheology further reveals how these crosslinkers govern strain-stiffening or softening behavior: passive crosslinkers (such as filamin or fascin) promote strain stiffening through filament alignment and bundling [11, 49, 50], while active crosslinkers like myosin II generate contractile prestress, enhancing strain stiffening or, under certain conditions, inducing network fluidization [21, 15, 36, 51, 52].

Despite extensive studies of actin networks crosslinked by passive proteins or active motors, a key unresolved question is how systematic variation in myosin content quantitatively modulates mechanical properties across both linear and non-linear regimes. While prior work has established that myosin can fluidize or stiffen networks depending on ATP conditions or crosslinker context, the systematic effect of myosin concentration on viscoelastic moduli, stress relaxation dynamics, and strain-stiffening behavior remains poorly understood. Moreover, few studies have directly linked motor-induced structural remodeling, as visualized through confocal microscopy, to corresponding changes in microscale mechanics measured by high-resolution active microrheology. Understanding this coupling is essential for elucidating how cells tune their cytoskeletal architecture and mechanics through differential motor engagement.

Here, we study the mechanical behavior of reconstituted actomyosin networks, focusing on how varying the myosin-to-actin molar ratio (*R*_MA_) tunes their structural organization and mechanical performance. By combining confocal fluorescence microscopy and optical tweezers microrheology, we probe the linear viscoelastic moduli, nonlinear strain-stiffening responses, and post-deformation relaxation dynamics of actomyosin networks. This work directly investigates how active motor content modulates network stiffness, energy dissipation, and stress retention, providing mechanistic insights that bridge simplified in vitro systems with the complex composite mechanics of living cells.

## 2 MATERIALS AND METHODS

### 2.1 Sample Preparation

Rabbit skeletal actin (Cat.# AKL99), chicken skeletal myosin II (Cat.# MY02), and Rhodamine Phalloidin (Cat.# PHDR1) were obtained from Cytoskeleton, Inc. Actin was stored at −80^°^C in G-buffer [2 mM Tris (pH 8.0), 0.5 mM DTT, 0.1 mM CaCl_2_, 0.2 mM ATP]. Myosin II was stored at −80^°^C in storage buffer [600 mM KCl, 25 mM KPO_4_, 10 mM EDTA, 1 mM DTT], and rhodamine phalloidin was kept at −20^°^C in 100% methanol. Prior to use, myosin II was dialyzed against 300 mM KCl for 4 hours using a 10 kDa MWCO microdialysis plate (ThermoFisher), with buffer exchange at 2 hours, and kept on ice thereafter.

All samples were prepared in thin rectangular chambers constructed by sealing a glass coverslip to a glass slide with double-sided tape. Actomyosin networks were reconstituted by polymerizing actin in the presence of myosin II at varying myosin-to-actin molar ratios (*R*_MA_) ranging from 0 to 0.08, while keeping the total actin concentration and *α*-actinin to actin ratio fixed at 5.8 *μ*M and 0.05, respectively. Actin monomers were labeled at a 1:10 ratio (labeled: unlabeled) using rhodamine phalloidin and polymerized in F-buffer [10 mM Imidazole (pH 7.0), 50 mM KCl, 1 mM MgCl_2_, 1 mM EGTA, 0.2 mM ATP] for 1 hour at room temperature. This concentration corresponds to a nominal mesh size of *ζ* ≈ 0.6 *μ*m, estimated from 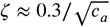 for *c*_*a*_ = 0.25 mg/ml [53, 54, 55]. The ATP concentration was maintained at 1 mM for *R*_MA_ *>* 0 to support myosin ATPase activity. Polystyrene microspheres (Bangs Laboratories, Inc., radius *a* = 2.1 *μ*m) were added at low volume fractions for microrheology. Beads were selected to be significantly larger than the mesh size to ensure they probed bulk network properties [56, 57, 58].

### 2.2 Confocal Fluorescence Microscopy

To visualize the network architecture, actin filaments were labeled with Acti-Stain 555 Rhodamine Phalloidin and imaged using a Leica TCS SP5 laser scanning confocal microscope equipped with a 60× 1.4 NA oil immersion objective. An oxygen scavenging system (4.5 *μ*g/mL glucose, 0.005% *β*-mercaptoethanol, 4.3 *μ*g/mL glucose oxidase, and 0.7 *μ*g/mL catalase) was used to suppress photobleaching. Three-dimensional *z*-stack images (42 slices at 0.5 *μ*m spacing) were collected at 512×512 pixel resolution over a 246 × 246 *μ*m field of view for each *R*_MA_. In parallel, time-lapse sequences (187 frames at 1 fps over 3 minutes) were acquired from the same field of view for all conditions. The *α*-actinin crosslinker concentration was fixed at 0.05 to ensure consistent connectivity across conditions.

### 2.3 Spatial Image Correlation Analysis

To quantify network organization, spatial image autocorrelation analysis was performed on both *z*-stacks and time-lapse projections using custom Python scripts [59, 60]. The radial correlation function *g*(*r*) was computed from the fluorescence intensity field *I* (*r*) using: *g*(*r*) = *F* ^−1^(| *F* (*I* (*r*)) |^2^)/[*I* (*r*)]^2^ where *F* and *F* ^−1^ denote the fast Fourier and inverse Fourier transforms, respectively. The structural correlation length *ξ* was extracted by fitting *g*(*r*) to an exponential decay function *g*(*r*) = *g*_0_ exp(−*r*/*ξ*) over the range *r* ≤ 5 *μ*m, following previous protocols used to quantify filament bundling and coarsening in cytoskeletal networks [28, 61].

### 2.4 Optical Tweezers Microrheology

All microrheology experiments were conducted using a custom-built optical tweezers (OT) system based on an Olympus IX73 fluorescence microscope. A 1064 nm Nd:YAG fiber laser (BKtel, RPMC Lasers, Inc.) was focused through a 63×, 1.42 NA oil immersion objective (UPLXAPO60XO, Olympus) to trap individual polystyrene microspheres (radius *a* = 2.1 *μ*m) embedded in the actomyosin network. Trap stiffness *κ* was calibrated independently via both passive equipartition and active Stokes drag methods [62, 63, 26]. The force exerted on the bead was inferred from laser deflection recorded by a position-sensing detector (PSD; PSM2-10Q, ON-TRAK Photonics) coupled with amplification via an OT-301 amplifier.

### 2.5 Linear Microrheology

To measure linear viscoelastic properties, the sample stage was held fixed and thermal fluctuations of the trapped bead were recorded for 130 s at 20 kHz across 25 independent beads at distinct locations (see Fig. 1(B)). Frequency-dependent storage and loss moduli, *G*^′^(*ω*) and *G*^′′^(*ω*), were extracted via the generalized Stokes–Einstein relation (GSER), using the Fourier-transformed normalized mean squared displacement 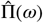 [64]. The time-domain normalized MSD was calculated as: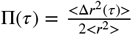, and its frequency-domain counterpart 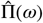 was obtained using oversampled PCHIP interpolation in MATLAB and the relation [64]:

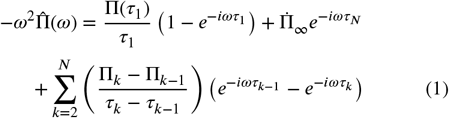

where 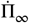 is the slope of Π(*τ*) extrapolated to infinite time, *τ* is the lag time, *τ*_*N*_ is the *N*^*th*^ lag time, and 1 and N represent the first and last point of the oversampled Π(*τ*). The complex modulus *G*^*\**^(*ω*) was determined from 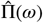 as:

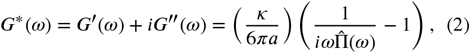

where *a* is the bead radius and *κ* is the trap stiffness. From *G*^*\**^(*ω*), we computed the complex viscosity *η*^*\**^(*ω*) = |*G*^*\**^(*ω*) |/*ω* and the loss tangent tan *δ* = *G*^′′^/*G*^′^. The data presented in Fig. 3 represent ensemble averages over 25 trials, with error bars corresponding to the standard error of the mean.

**FIGURE 1.**
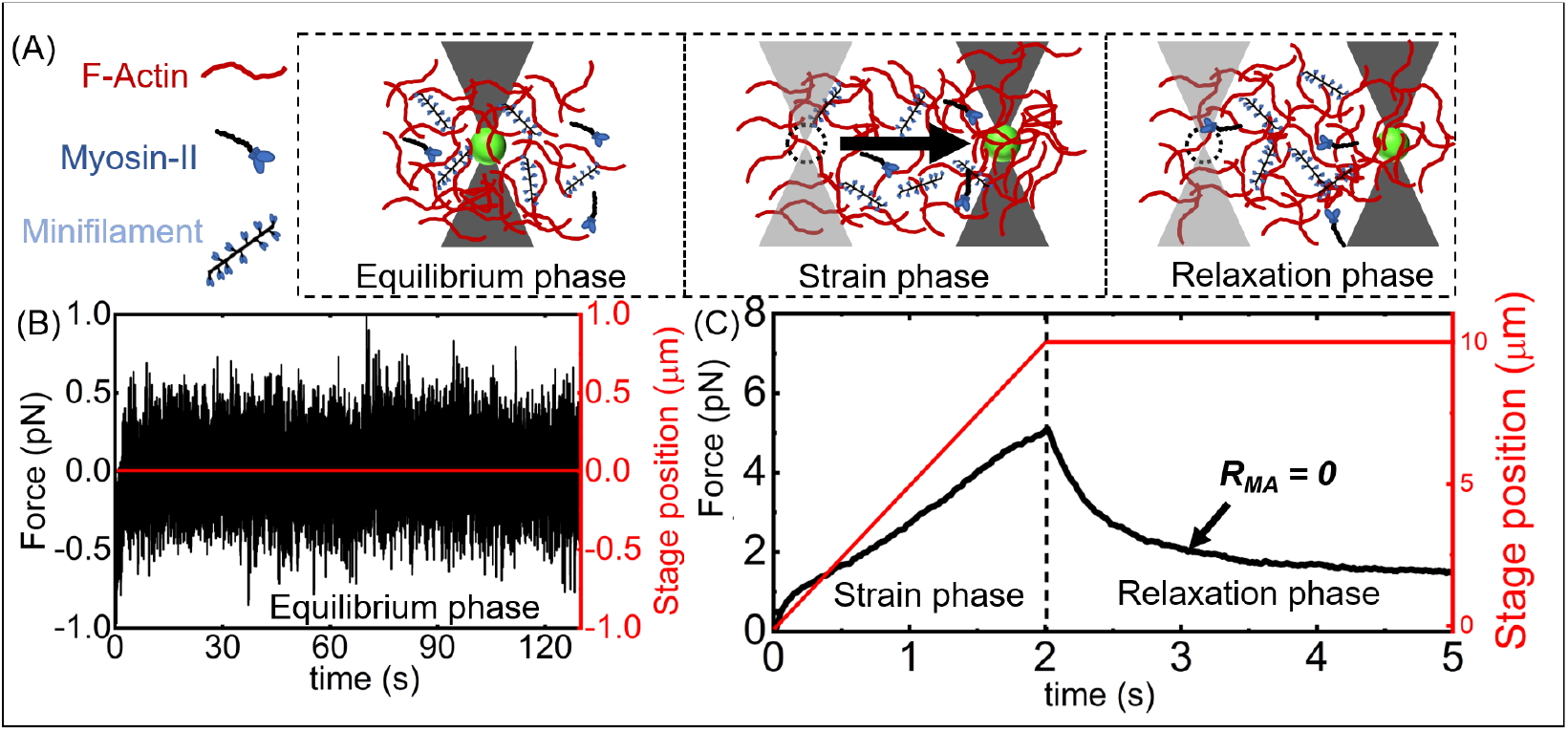
Optical tweezers microrheology of actomyosin networks. (A) Schematic of actin network (red) cross-linked by myosin II minifilament (blue) at three measurement phases: equilibrium (probe particle (green) held stationary), strain (middle, trapped probe moves 10 *μ*m at *v* = 5 *μ*m/s), and relaxation (no trap movement, probe remains trapped while network recoils). *α*-actinin crosslinkers are present but not depicted. (B) In the equilibrium phase measurement, the stage position (shown in red) is held fixed while the thermal force fluctuations (depicted in black) are measured for a duration of 130 seconds. This allows for the extraction of the linear viscoelastic moduli, *G*^′^(*ω*) and *G*^′′^(*ω*). (C) In the nonlinear measurement, we record the force during both the strain and relaxation phases. During the strain phase, the probe particle is pulled 10 *μ*m by a piezoelectric stage, indicated in red. In the relaxation phase, the position of the bead is held constant while we measure the force. The sample data shown in (B) and (C) are for the *R*_MA_ = 0 condition. In C, only a portion of the relaxation force is shown, while in the experiment, we collect data for 15 seconds.

**FIGURE 2.**
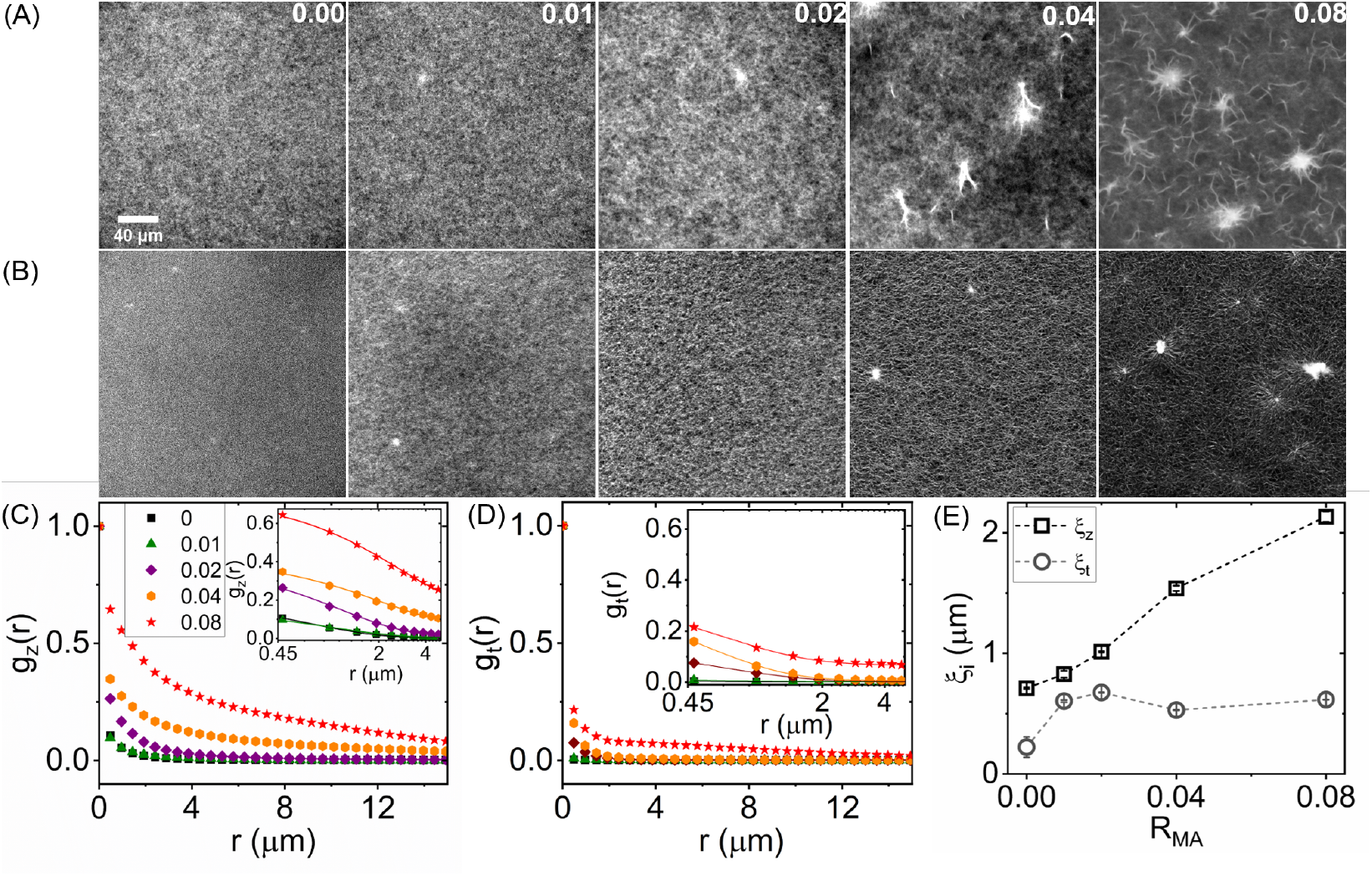
Increasing the myosin-to-actin molar ratio *R*_MA_ at a constant total actin concentration (*c*_*a*_ = 5.8 *μ*M) and fixed *α*-actinin crosslinker ratio (0.05) induces pronounced bundling and structural heterogeneity in reconstituted actomyosin networks. Shown are mean intensity projections of (A) *z*-stacks (42 slices at 0.5 *μ*m spacing) and (B) time-lapse recordings (187 frames acquired at 1 fps over 3 minutes), captured using a Leica TCS SP5 confocal microscope with a 60×, 1.4 NA objective. (C, D) Radial spatial autocorrelation functions *g*_z_(*r*) and *g*_t_ (*r*), computed from the *z*-stacks and time-lapse projections, respectively, quantify changes in network organization with increasing *R*_MA_. Insets show *g*(*r*) on a log-linear scale, highlighting exponential decay behavior. (E) Structural correlation lengths *ξ*_z_ and *ξ*_t_ were extracted by fitting 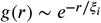 where *ξ*_*i*_ ∈ (*ξ*_z_, *ξ*_t_) over distances *r* ≤ 5 *μ*m, capturing short-range coherence. Error bars denote standard errors across 42 *z*-planes and time points. Scale bars: 40 *μ*m.

**FIGURE 3.**
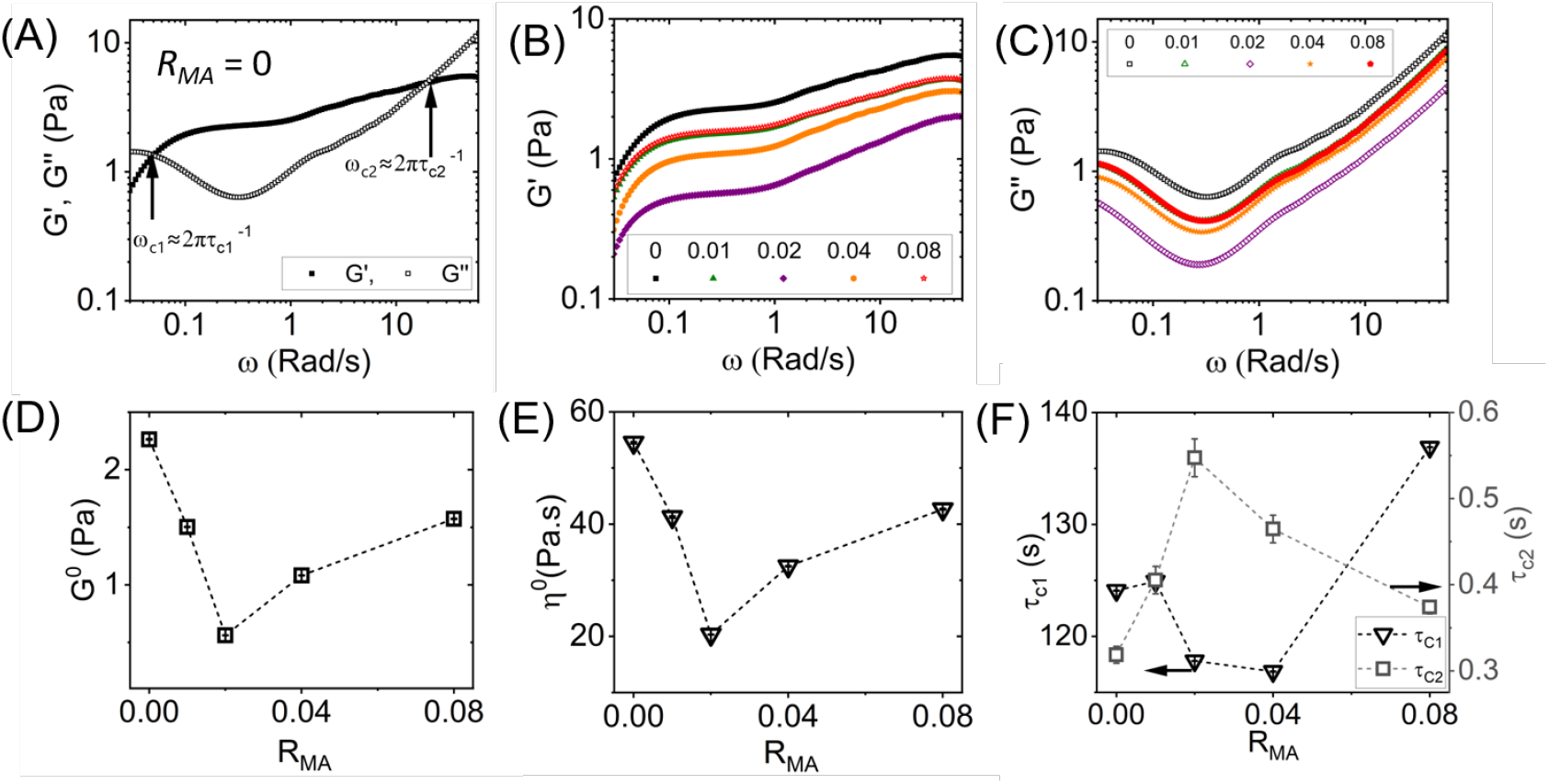
Linear microrheological properties of actomyosin networks exhibit non-monotonic dependence on the myosin-to-actin molar ratio *R*_MA_. (A–C) Frequency-dependent elastic (*G*^′^(*ω*), filled symbols) and viscous (*G*^′′^(*ω*), open symbols) moduli for (A) F-actin networks in the absence of myosin (*R*_MA_ = 0), compared to (B, C) actomyosin networks with increasing myosin content (*R*_MA_ = 0.01–0.08). (D) The quasi-plateau modulus *G*^0^, and (E) the zero-shear viscosity *η*^0^, both exhibit a non-monotonic dependence on *R*_MA_, reaching minima near *R*_MA_ = 0.02. (F) Long (left y-axis, indicated by the black left-pointing arrow) and short (right y-axis, indicated by the black right-pointing arrow) relaxation times 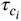 computed from the crossover frequencies 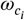 at which *G*^′^(*ω*) = *G*^′′^(*ω*), using the relation 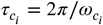. Representative slow and fast crossover frequencies 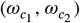, corresponding to long and short relaxation times 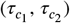, are indicated in panel (A).

### 2.6 Nonlinear Microrheology

To probe nonlinear mechanical responses, trapped microspheres were displaced through the network by 10 *μ*m at a constant speed of 5 *μ*m/s using a nanopositioning piezoelectric stage (PDQ-250, Mad City Laboratories) to move the sample chamber relative to the fixed trap. This strain phase (2 s) was followed by a 15 s relaxation phase during which the trap was held stationary and the network recoiled. Laser deflection was recorded throughout at 20 kHz (see Fig. 1C). The resulting force-displacement and relaxation curves, shown in Figs. 4 and 5, represent averages over two sets of 50 trials, each with unique bead locations. The applied strain rate was 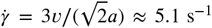, a regime previously shown to exceed the threshold for onset of nonlinear response in actin networks [65]. Custom LabVIEW software was used for instrument control and data acquisition, while all post-processing was performed in MATLAB. Error bars on non-linear results were determined by bootstrapping over 1000 resampled subsets.

**FIGURE 4.**
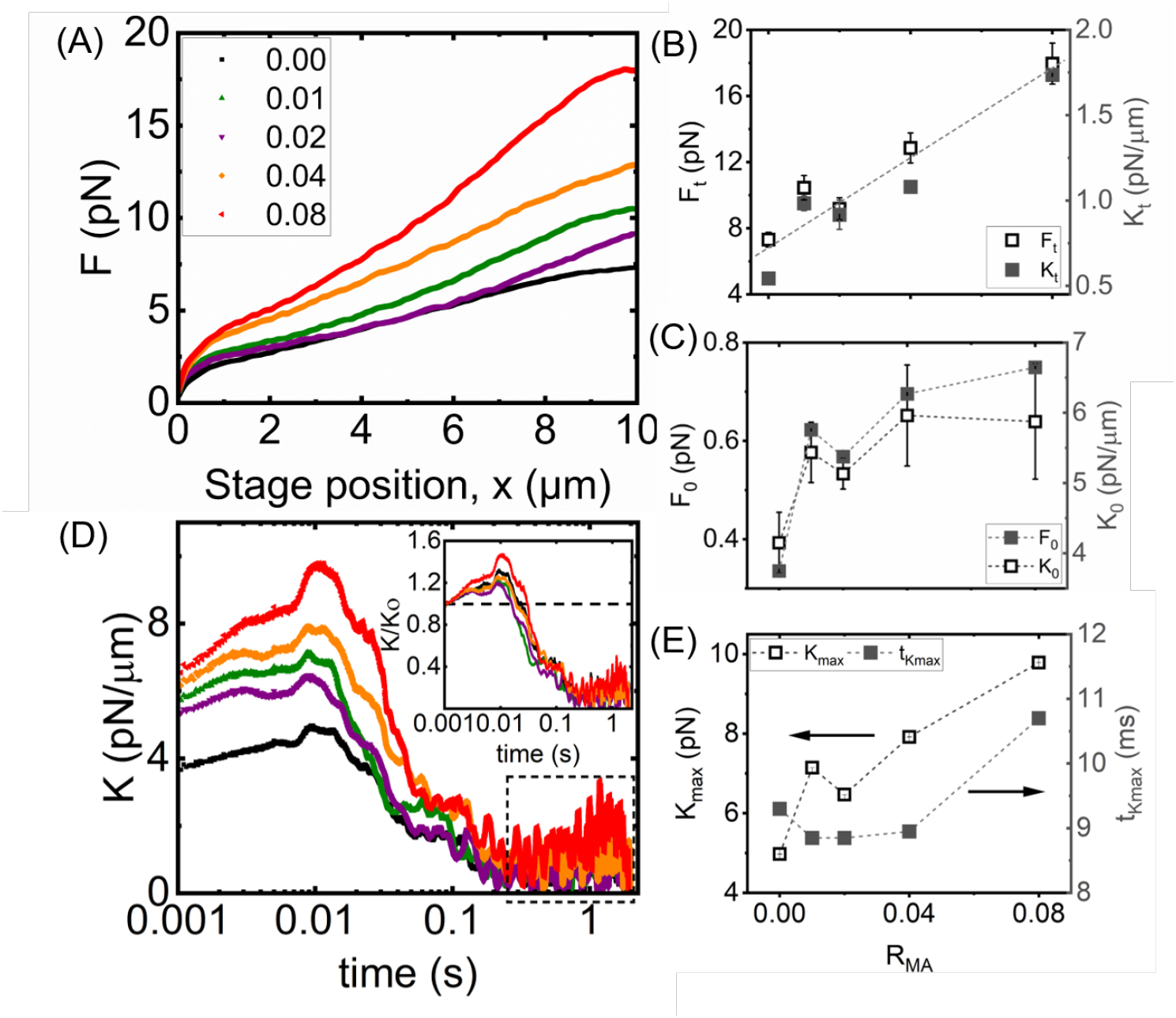
Nonlinear microscale force response of actomyosin networks during the strain phase. (A) Force response as a microsphere is pulled through the network at a constant speed of 5 *μ*m/s, plotted versus stage displacement *x*. (B) Terminal force *F*_t_ (left y-axis, open black squares) and terminal stiffness *K*_t_ (right y-axis, filled grey squares) as functions of myosin-to-actin molar ratio *R*_MA_, measured at the end of the strain (*x* = 10 *μ*m). (C) Initial force *F*_0_ (left y-axis, open black squares) and initial stiffness *K*_0_ (right y-axis, filled grey squares), determined at the onset of strain (*x* = 0), showing the increasing yield threshold and prestress with *R*_MA_. (D) Time evolution of the effective differential modulus, *K*(*t*) = *dF* /*dx*, computed from the average *F* (*x*) curves shown in (A). Inset: Normalized modulus *K*(*t*)/*K*_0_, highlighting the degree of strain stiffening (*K*/*K*_0_ *>* 1) across motor concentrations. (E) Maximum differential modulus *K*_max_ (left y-axis, open black squares, indicated by the black left-pointing arrow) and the corresponding time to peak stiffening 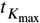 (right y-axis, filled grey squares, indicated by the black right-pointing arrow) as functions of *R*_MA_. While *K*_max_ increases monotonically with motor content, 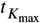 exhibits a non-monotonic trend, decreasing at low *R*_MA_ and plateauing at higher values. Error bars represent the standard error across individual trials contributing to the average curves in (A), as shown in SI Fig. S2. The dotted lines are guides to the eye.

**FIGURE 5.**
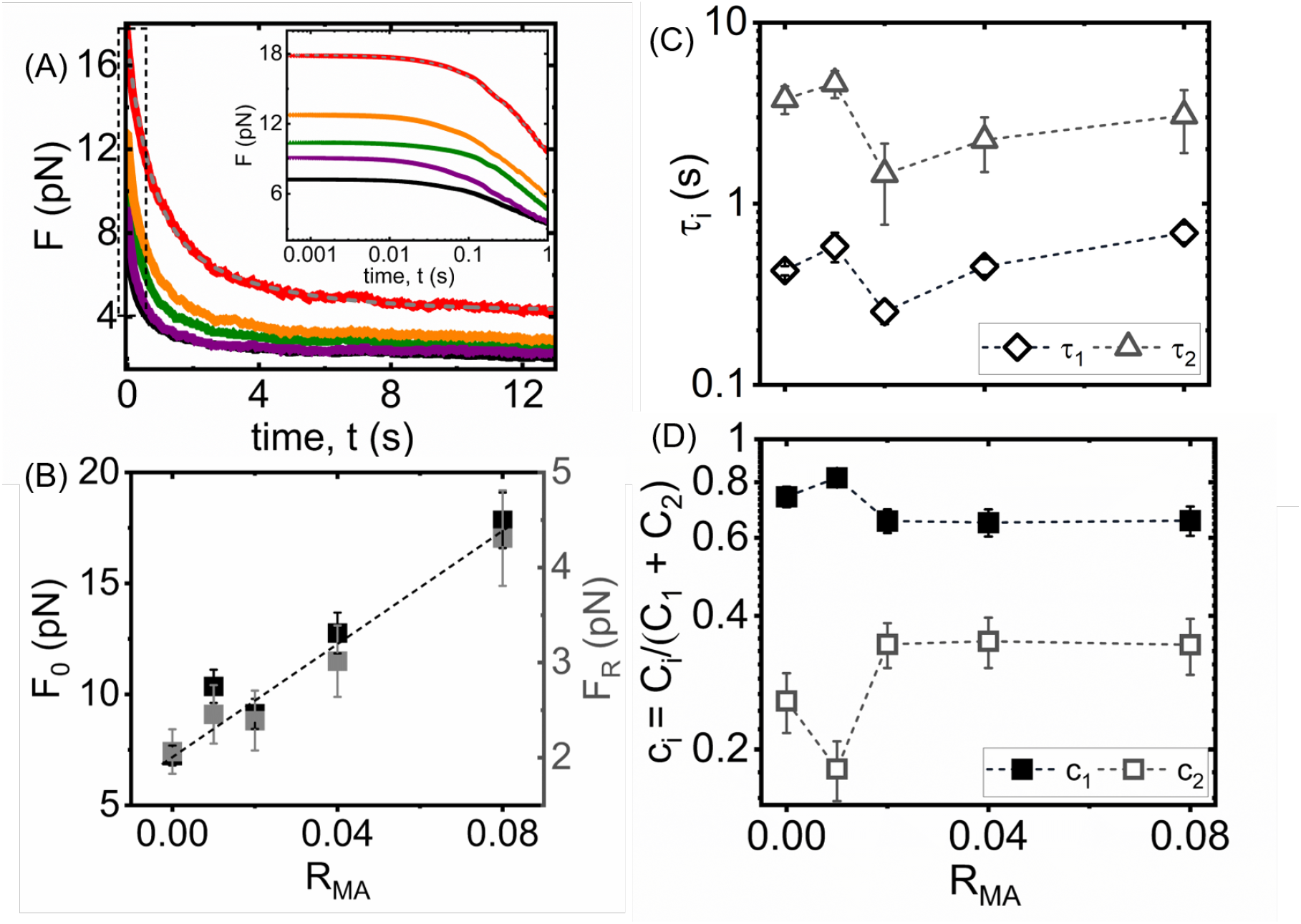
Relaxation dynamics of induced force during the post-strain relaxation phase. (A) Relaxation force as a function of time following the cessation of bead motion, measured after probe particles were pulled through actomyosin networks with varying myosin-to-actin molar ratios *R*_MA_ (colors as in Fig. 4A) at a constant speed of 5 *μ*m/s. (Inset) Zoom-in of small t region denoted by dashed box to show the dependence of the initial force *F*_0_ on *R*_MA_. (B) Initial force at the start of the relaxation phase (left y-axis, filled black squares) and the average terminal (residual) force *F*_R_ (right y-axis, filled grey squares), plotted as functions of *R*_MA_. (C) Fast (*τ*_1_, open diamonds) and slow (*τ*_2_, open squares) relaxation times extracted from fits to the relaxation curves in (A), as functions of *R*_MA_. Relaxation dynamics were fit using a biexponential model: *F* (*t*) = *C*_1_ exp(−*t*/*τ*_1_) + *C*_2_ exp(−*t*/*τ*_2_) + *F*_R_. (D) Corresponding fractional contributions *c*_1_ = *C*_1_/(*C*_1_ + *C*_2_) and *c*_2_ = *C*_2_/(*C*_1_ + *C*_2_) derived from the same fits, plotted as functions of *R*_MA_. Error bars represent the standard error across individual trials used to compute the averaged relaxation curves in (A).

## 3 RESULTS

In this study, we investigated the structural and mechanical properties of actomyosin networks reconstituted in vitro by varying the molar ratio of myosin II to actin (*R*_MA_) from 0 to 0.08, while keeping the total actin concentration and *α*-actinin crosslinker density constant at *c*_*a*_ = 5.8 *μ*M and *R*_*αA*_ = 0.05, respectively.

### 3.1 Network morphological transitions with increasing motor content

To understand how myosin modulates the structural organization of actin networks, we reconstituted actomyosin networks with varying myosin-to-actin molar ratios (*R*_MA_) and performed laser scanning confocal microscopy. Representative average-intensity *z*-stack projections (Fig. 2A) and time-lapse projections (Fig. 2B) reveal a striking progression of structural reorganization as *R*_MA_ increases. At *R*_MA_ = 0, the F-actin network appears isotropic and homogeneous, consistent with a weakly entangled and crosslinked mesh of semiflexible filaments [66, 67]. Introducing low myosin concentrations (*R*_MA_ = 0.01 − 0.02) leads to the emergence of localized filament density variations, suggesting the onset of contractile activity and filament rearrangement. At higher motor content (*R*_MA_ = 0.04 and 0.08), the networks transform into spatially heterogeneous morphologies marked by thick bundles, aster-like foci, and large voids–features that reflect motor-driven contractile aggregation and active network coarsening [33, 20, 68].

To quantitatively characterize these morphological transitions, we computed spatial autocorrelation functions *g*_z_(*r*) and *g*_t_ (*r*) from the grayscale fluorescence intensity fields of the *z*-stacks and time-lapse projections, respectively (Figs. 2C,D). These functions capture the characteristic length scale over which filament density is spatially correlated [59, 60]. At low *R*_MA_, both *g*_z_(*r*) and *g*_t_ (*r*) decay rapidly with distance, consistent with a uniform fine-scale filament distribution. With increasing *R*_MA_, the decay becomes slower and broader, indicating the emergence of longer-range structural coherence and bundling. Insets in Figs. 2C and D show that the curvature of *g*(*r*) at short distances also steepens with motor content, reflecting enhanced local order.

Fitting the correlation functions to an exponential decay, *g*(*r*) ∼ *e*^−*r*/*ξ*^, yielded the correlation lengths *ξ*_z_ (from *z*-stacks) and *ξ*_t_ (from time projections) (Fig. 2E). These length scales provide a direct measure of spatial coherence or cluster size across the network [59, 60]. To minimize the influence of non-exponential tails at large distances—often caused by flat-field noise, as noted in previous studies [28, 61]–we restricted the fitting range to *r* ≤ 5 *μ*m. Notably, *ξ*_z_ increases approximately monotonically with *R*_MA_. This monotonic increase confirms that myosin motors actively drive actin coarsening by generating contractile stress that pulls filaments into mechanically cohesive domains. In contrast, *ξ*_t_ displays a non-monotonic trend: it initially increases, peaks around *R*_MA_ = 0.02, and then declines at higher motor levels. The divergence between *ξ*_z_ and *ξ*_t_ at high *R*_MA_ highlights the complex interplay between motor-driven contractility and network remodeling kinetics. The slight saturation of *ξ*_*t*_ at high *R*_MA_ likely results from the formation of percolated contractile clusters that begin to dominate the network architecture.

### 3.2 Linear microrheology reveals fluidization–reinforcement crossover

We utilized optical tweezers-based active microrheology to quantify the frequency-dependent elastic modulus *G*^′^(*ω*) and viscous modulus *G*^′′^(*ω*) of reconstituted actomyosin networks across varying myosin-to-actin molar ratios *R*_MA_ (see Methods). All samples exhibited characteristic viscoelastic solid behavior: the elastic modulus *G*^′^(*ω*) showed weak frequency dependence over intermediate frequencies, defining a quasi-plateau modulus, while the viscous modulus *G*^′′^(*ω*) dominated at low frequencies, indicative of fluid-like stress dissipation. As frequency increased, *G*^′′^(*ω*) decreased and eventually intersected *G*^′^(*ω*) at distinct crossover frequencies 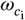, marking transitions between relaxation regimes (Fig. 3A). At even higher frequencies beyond the second crossover point, *G*^′′^(*ω*) once again exceeded *G*^′^(*ω*), reflecting dominant viscous dissipation arising from fast, local-scale processes such as filament bending, transient crosslinker dynamics, and solvent drag–features commonly observed in semiflexible polymer networks [22].

The addition of myosin had a pronounced non-monotonic effect on the linear viscoelastic moduli of the network. To characterize this behavior, we quantified the plateau modulus *G*^0^, defined as the elastic modulus *G*^′^ evaluated at the frequency where the loss tangent, tan *δ* = *G*^′′^/*G*^′^ reaches its minimum (SI Fig. S1B) [69, 70]. In the absence of myosin (*R*_MA_ = 0), *G*^0^ was relatively high–on the order of several Pascals–reflecting the elastic response of a densely entangled and transiently crosslinked actin scaffold. Upon the introduction of low levels of myosin (*R*_MA_ = 0.01–0.02), the network exhibits clear mechanical softening. As shown in Fig. 3B, the storage modulus *G*^′^(*ω*) decreases across the entire frequency spectrum relative to the myosin-free case. This reduction is particularly notable at low frequencies, where the network’s elastic response is most sensitive to structural integrity. The plateau modulus *G*^0^, shown in Fig. 3D, decreases by approximately 40–80% compared to the passive network, confirming a substantial loss in network stiffness. The viscous modulus *G*^′′^ followed a similar non-monotonic trend, as shown in Fig. 3C, initially decreasing at low motor concentrations, then rising at higher *R*_MA_. These results suggest that low levels of motor activity disrupt filament connectivity or prestrain without yet organizing the network into contractile bundles, leading to an overall softening effect.

However, as the myosin content increased further (*R*_MA_ = 0.04–0.08), the network exhibited partial mechanical recovery, as evidenced by a rise in the plateau modulus *G*^0^ relative to the intermediate minimum likely through enhanced bundling or contractile restructuring. Nevertheless, *G*^0^ remained comparable to or slightly below the value observed in the absence of myosin (Fig. 3D). This non-monotonic dependence of *G*^0^ on *R*_MA_–a sharp decrease at intermediate *R*_MA_ followed by a partial rebound at higher motor concentrations–suggests an initial fluidization of the network, succeeded by reinforcement due to increased internal dissipation and motor-induced restructuring.

A similar non-monotonic trend was observed in the zero-shear viscosity, *η*^0^, determined from the low-frequency limit of the dynamic viscosity *η*^*\**^(*ω*) (see SI Fig. S1C). For the motor-free actin network (*R*_MA_ = 0), *η*^0^ was relatively high, around ∼55 Pa·s, reflecting long-lived entanglements and crosslinks that slow stress relaxation. As *R*_MA_ increased to 0.02, *η*^0^ dropped sharply to approximately ∼20 Pa·s, consistent with motor-induced fluidization: the active disruption of network connectivity accelerates stress dissipation and reduces the effective viscosity [15, 37]. However, at the highest tested myosin concentration (*R*_MA_ = 0.08), *η*^0^ rose again to nearly ∼43 Pa·s, indicating that the network had reorganized into a more solid-like, contractile state capable of storing long-lived internal stresses [71, 20]. This parallel modulation of both *G*^0^ and *η*^0^ reinforces the view that actomyosin activity enables dynamic tuning of viscoelastic behavior through a balance of structural disruption and contractile reorganization.

To further explore how these structural transitions shape relaxation dynamics across timescales, we examined the characteristic relaxation times 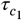 (slow mode) and 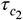 (fast mode), extracted from the crossover frequencies 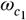 and 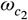, defined by the intersections of *G*^′^(*ω*) and *G*^′′^(*ω*) at low and high frequencies (SI Fig. S1A), respectively, using the relation 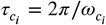. Notably, 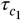 exhibited a modest but non-negligible non-monotonic trend: it decreased from ∼125 s at *R*_MA_ = 0 to a minimum of ∼115 s near *R*_MA_ = 0.02, followed by a progressive increase to ∼137 s at *R*_MA_ = 0.08. This shallow minimum suggests that moderate motor activity transiently accelerates large-scale structural relaxation, likely by disrupting filamentous connectivity. In contrast, 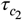 displayed a more pronounced non-monotonic response: starting at ∼0.3 s without myosin, peaking at ∼0.6 s near *R*_MA_ = 0.02, and decreasing to ∼0.35 s at *R*_MA_ = 0.08. This pattern indicates a dynamic interplay between motor-induced remodeling, which initially slows local relaxation dynamics, and the formation of an increasingly rigid network at higher motor content.

Overall, these linear measurements reveal that introducing low-to-intermediate levels of myosin initially fluidizes the actin network, reducing *η*^0^ and altering both slow and fast relaxation dynamics. However, at high motor density, the system transitions into an actively reinforced state, with elevated viscosity and enhanced stress storage [37, 15]. This non-monotonic crossover behavior underscores the rich mechanical tunability of actomyosin networks and highlights the capacity of molecular motors to modulate both linear viscoelasticity and multiscale stress relaxation across compositional regimes.

### 3.3 Nonlinear microrheology reveals tunable stiffening and stress retention

To investigate how motor activity modulates cytoskeletal mechanics in the nonlinear regime, we performed constant-speed strain experiments using optical tweezers. A microsphere embedded in the network was displaced by 10 *μ*m at a speed of 5 *μ*m/s, and the resulting force *F* on the bead was recorded as a function of stage position *x*. As shown in Fig. 4A, the force– displacement curves *F* (*x*) exhibit progressively steeper slopes and higher terminal forces with increasing myosin content, indicating enhanced mechanical resistance. In the passive actin-only network (*R*_MA_ = 0), *F* (*x*) increased gradually, reaching a terminal force of approximately *F*_t_ ∼ 5–6 pN at *x* = 10 *μ*m, consistent with low resistance to deformation. Upon introduction of myosin, the response was markedly amplified: at *R*_MA_ = 0.01, *F*_t_ nearly doubled to ∼10 pN, and at the highest motor concentration tested (*R*_MA_ = 0.08), it rose to ∼15–20 pN, reflecting substantial contractile reinforcement. This trend is quantified in Fig. 4B (left axis), where the terminal force *F*_t_ increases monotonically with *R*_MA_. The enhanced *F*_t_ values in myosin-rich networks reflect increased prestress and filament bundling, underscoring the capacity of motor activity to reinforce network connectivity and mechanical resilience. We next considered the initial force *F*_0_, defined at the onset of strain (*x* = 0), which primarily reflects prestrain tension built up in the network before active displacement begins. As shown in Fig. 4C (left axis), *F*_0_ also rises with *R*_MA_, indicating that higher motor content leads to more prestressed initial configurations. This increase in *F*_0_ suggests that myosin generates internal contractile stresses that precondition the network even prior to externally applied load.

To further dissect the evolution of network mechanics under strain, we computed the effective differential modulus *K*(*t*) = *dF* /*dx* from the force–displacement data in Fig. 4A. As shown in Fig. 4D, all networks—including the passive actin-only system—exhibited some degree of strain stiffening (*dK*/*dt >* 0) at short times (*t <* 0.01 s). Notably, even the passive network shows *K*(*t*)/*K*_0_ *>* 1 (inset), indicating intrinsic nonlinear elasticity. With increasing *R*_MA_, networks not only achieve greater peak stiffness (*K*_max_), but also stiffen more rapidly, implying faster recruitment of load-bearing filament structures. These trends highlight that myosin activity amplifies and accelerates nonlinear mechanical reinforcement during strain. Following an initial stiffening to a peak value *K*_max_, the networks transition into strain softening (*dK*/*dt <* 0) followed by a terminal regime characterized by low, nearly viscous stiffness values *K*_*t*_, which show minimal dependence on *x* (Figure 4B, right) over a timescale of ≈ 0.5 s. Complementing this dynamic analysis, we evaluated the terminal stiffness *K*_t_, obtained by averaging the K values shown in the dotted box in Fig. 4D. As illustrated in Fig. 4B (right axis), the terminal stiffness *K*_t_ increases steadily with *R*_MA_. This cincrease is closely tracking the rise in terminal force *F*_t_. Such a trend indicates that as motor content increases, the network becomes progressively more resistant to deformation, reflecting the development of load-bearing structures during strain.

In contrast, the initial stiffness, *K*_0_, measured at the onset of strain (*t* = 0), reveals a non-monotonic dependence on motor concentration. As shown in Fig. 4C (right axis), *K*_0_ rises sharply with low levels of myosin, peaks near *R*_MA_ ≈ 0.02, and then declines at higher motor concentrations. This non-monotonic trend contrasts with the monotonic increase in *F*_t_ and *K*_t_, and suggests that short-time elasticity arises from a distinct balance of filament connectivity and dynamic crosslinking. These observations support the emergence of a transiently optimized elastic state at intermediate motor densities [15, 68, 37]. Notably, the initial force *F*_0_ (Fig. 4C, left axis) follows a similar trend to *K*_0_ (Fig. 4C, right axis), further indicating that both early-time elastic force generation and stiffness are governed by the same underlying structural transitions within the network.

To further understand how strain-induced stiffening evolves over time and with motor content, we analyzed the maximum differential stiffness *K*_max_ and the time to reach peak stiffness 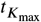, as shown in Fig. 4E. With increasing *R*_MA_, *K*_max_ exhibits a clear monotonic rise, indicating that myosin activity enhances the extent of stiffening during deformation. This trend supports the interpretation that elevated motor levels promote long-range force percolation and active network reinforcement. In contrast, 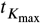 shows a non-monotonic dependence on *R*_MA_: it decreases slightly from ∼9.3 ms at low *R*_MA_ to a minimum near *R*_MA_ ≈ 0.01, and then plateaus or slightly increases. This biphasic behavior suggests that while moderate motor levels accelerate stress buildup and network reorganization, further increases in motor density may saturate the kinetics of contraction or filament alignment, limiting additional gains in stiffening speed.

### 3.4 Stress relaxation follows multi-timescale decay with residual force

Upon completion of bead displacement, the trap position was held constant and the force exerted on the bead was recorded during network relaxation. The resulting force-relaxation curves *F* (*t*), shown in Fig. 5A, were well described by a bi-exponential decay with a residual offset: 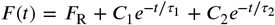, where *F*_R_ represents the long-time residual force, *τ*_1_ and *τ*_2_ are fast and slow relaxation time constants, and *C*_1_, *C*_2_ are the corresponding amplitudes. This functional form captured the data across all motor concentrations, indicating that stress relaxation proceeds via two distinct timescales.

We first examined the force at the onset of the relaxation phase, *F*_0_, which increases systematically with the myosin-to-actin ratio *R*_MA_ (Fig. 5B, left axis, black squares). This monotonic rise is consistent with enhanced contractile buildup during the preceding strain phase and mirrors the trends observed for the peak force and terminal stiffness (Fig. 4B), reinforcing the role of myosin in generating internal prestress. To evaluate the extent of stress retention, we quantified the residual force *F*_R_ maintained at the end of the relaxation phase (Fig. 5B, right axis, grey squares). In the passive actin network (*R*_MA_ = 0), the force decayed to near-zero levels (*<* 2 pN), consistent with complete stress dissipation. However, *F*_R_ increased approximately linearly with motor content, reaching ∼ 5 pN at *R*_MA_ = 0.08. This partial force retention indicates that actomyosin networks, particularly at higher motor densities, develop long-lived, stress-bearing configurations that resist full relaxation. To quantify the degree of mechanical memory, we compared the residual force *F*_R_ to the terminal force *F*_t_ reached during the strain phase (Figs. 4B, 5B). In motor-free networks, only ∼10% of the force was retained after relaxation, indicative of predominantly viscous or plastic behavior. By contrast, networks with high myosin content retained nearly 30% of the applied force, suggesting substantial elastic recovery. These results align with previous findings in crosslinked microtubule systems [72] and actomyosin simulations [73], where 25–50% of induced deformation was reported to be elastically recoverable. The residual force observed here likely reflects motor-stabilized, crosslinked domains that maintain internal tension, highlighting the role of active processes in modulating mechanical memory and viscoelastic response.

To elucidate how motor activity modulates the temporal dynamics of stress dissipation, we analyzed the characteristic relaxation timescales extracted from force relaxation profiles. The extracted relaxation times *τ*_1_ and *τ*_2_ quantify the fast and slow modes of force dissipation within the actomyosin network (Fig. 5C). Both timescales exhibit nonmonotonic behavior as the myosin-to-actin ratio *R*_MA_ increases, revealing complex regulation of stress relaxation by motor activity. At the lowest motor level (*R*_MA_ = 0), the fast relaxation time *τ*_1_ is approximately 0.4–0.5 s. Interestingly, *τ*_1_ initially rises to ∼ 0.7 s at *R*_MA_ = 0.01, followed by a sharp decrease to ∼ 0.2 s at *R*_MA_ = 0.02, before gradually increasing again at higher motor concentrations, reaching ∼ 0.7 s at *R*_MA_ = 0.08. A similar trend is observed for the slow relaxation time *τ*_2_, which starts around 4 s at *R*_MA_ = 0, peaks near 6 s at *R*_MA_ = 0.01, and then declines to ∼ 3 s by *R*_MA_ = 0.08. Despite the small fluctuations, *τ*_2_ remains on average an order of magnitude longer than *τ*_1_ across all conditions (⟨*τ*_2_⟩ ≈ 3.04 ± 0.56 s; ⟨*τ*_1_⟩ ≈ 0.480 ± 0.074 s), highlighting distinct contributions of local and global processes to viscoelastic relaxation.

To shed further light on how motor activity influences stress relaxation dynamics in actomyosin networks, we quantified the relative contributions of the fast and slow relaxation modes by evaluating the fractional coefficients of the corresponding exponential decay terms: *c*_1_ = *C*_1_/(*C*_1_ + *C*_2_) and *c*_2_ = *C*_2_/(*C*_1_ +*C*_2_) (Fig. 5D). These quantities provide insight into how distinct physical mechanisms contribute to the overall relaxation response. Under passive conditions (*R*_MA_ = 0), network relaxation is dominated by the fast mode (*c*_1_ ≈ 0.7), consistent with rapid stress dissipation through filament-scale processes such as bending, recoil, or transverse fluctuations within confinement tubes [74, 75]. Upon the introduction of low myosin concentrations (*R*_MA_ = 0.01), *c*_1_ increases slightly to 0.8, suggesting enhanced local remodeling or filament mobility driven by active forces. As motor content increases further, *c*_1_ decreases to ∼0.6 at *R*_MA_ = 0.02 and then stabilizes, with an average value of ⟨*c*_1_⟩ ≈ 0.704 ± 0.033 across all conditions. The corresponding increase in the slow-mode fraction, ⟨*c*_2_⟩ ≈ 0.296 ± 0.033, reflects a growing contribution from slower relaxation processes, such as filament disengagement, crosslinker unbinding, or global network reorganization. This redistribution in relaxation weighting may result from increasing internal stresses or structural constraints that prolong stress decay. Notably, at low motor densities, the reduction in *c*_2_–indicating suppressed slow relaxation– coincides with the appearance of nonzero residual force *F*_R_ (Fig. 5B). This correlation suggests that diminished access to slow, long-timescale relaxation pathways limits the network’s ability to fully dissipate stress, resulting in greater stress retention. Meanwhile, the consistently high values of *c*_1_ across all *R*_MA_ values indicate that fast, filament-scale processes remain a dominant and robust mechanism for mechanical dissipation, even as active remodeling proceeds. These results likely illustrate how myosin motors modulate both local and global mechanical responses by altering the relative contributions of fast and slow relaxation dynamics.

### 3.5 Coupling between structure and mechanics

To elucidate the relationship between network structure and relaxation dynamics, we examined how the characteristic relaxation times 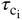 and *τ*_i_, derived from linear and nonlinear microrheology, respectively, vary as a function of the spatial correlation length *ξ*_*z*_ extracted from confocal *z*-stack projections.

As shown in Fig. 6A, among the linear regime relaxation times, the short time scale, 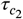 (gray squares), exhibits modest variation across the range of *ξ*_*z*_, rising slightly at intermediate values before returning to its baseline. In contrast, the long timescale, 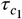 (black downward triangles), remains remarkably stable regardless of network structure. This insensitivity suggests that 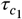 captures global, persistent stress-bearing modes that are decoupled from mesoscale remodeling.

**FIGURE 6.**
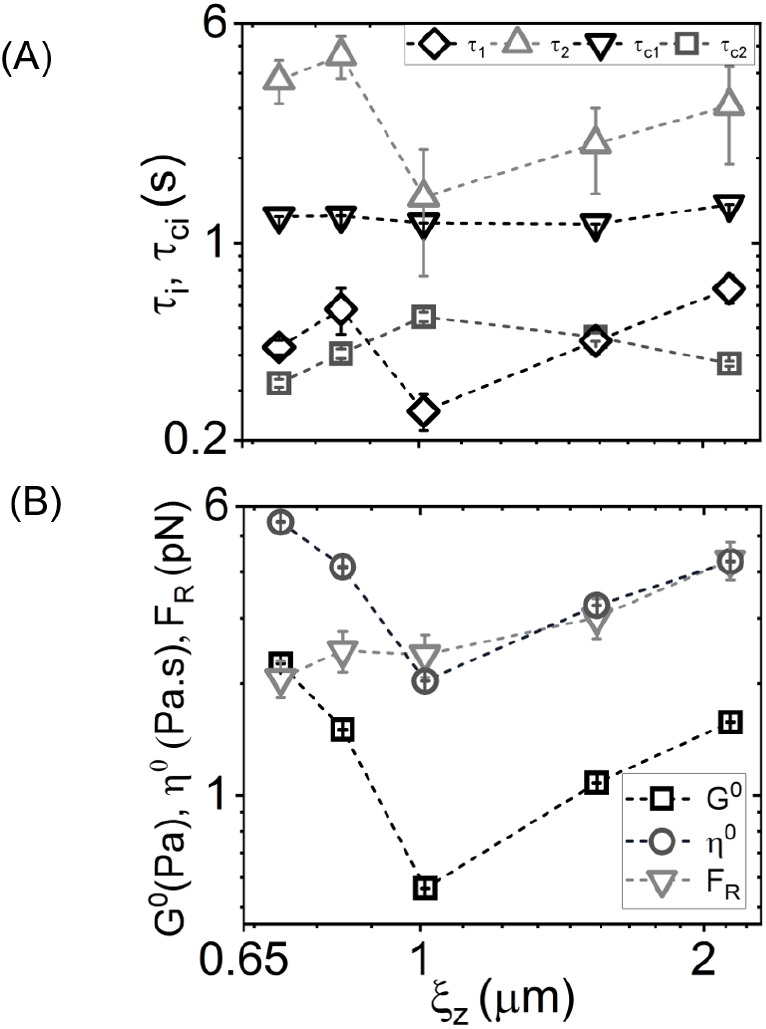
Structural–mechanical coupling in actomyosin networks. (A) Relaxation time constants *τ*_1_ (black diamonds), *τ*_2_ (gray triangles), 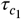 (black downward triangles), and 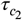 (gray squares), plotted as a function of structural correlation length *ξ*_*z*_. Both the short and long timescales *τ*_1_ and *τ*_2_ exhibit a pronounced non-monotonic trend, reaching a minimum near *ξ*_*z*_ ∼ 1 *μ*m, suggesting that intermediate levels of network coarsening facilitate more rapid local relaxation. In contrast, the longest timescale 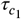 remains nearly constant, reflecting robust, architecture-independent stress-bearing modes while 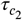 shows a shallow peak at intermediate *ξ*_*z*_, indicating modest slowing of intermediate-timescale stress dissipation in more coarsened architectures. (B) Mechanical parameters extracted from linear and nonlinear force relaxation measurements, including plateau modulus *G*^0^ (black squares), zero-shear viscosity *η*^0^ (gray circles), and residual force *F*_*R*_ (gray downward triangles), also plotted as a function of *ξ*_*z*_. *η*^0^ and 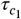 are scaled by 10 and 100, respectively, to fit in the panels. All three parameters increase with increasing *ξ*_*z*_ after the initial slight decrease. These trends indicate that actomyosin-driven structural coarsening enhances network stiffness, slows viscous relaxation, and promotes the buildup and retention of long-lived internal stress. Error bars represent the standard error across multiple measurements.

Turning to the nonlinear regime, the short-timescale relaxation time *τ*_1_ (black diamonds) exhibits a pronounced non-monotonic dependence on *ξ*_*z*_: it increases at low *ξ*_*z*_, reaches a minimum near *ξ*_*z*_ ∼ 1 *μ*m, and then rises again as coarsening progresses. This U-shaped behavior suggests that intermediate levels of motor-driven remodeling facilitate faster local relaxation–possibly due to transient disruption of filament connectivity–while highly coarsened networks reinforce local mechanics and slow stress dissipation. *τ*_2_ (gray triangles), the slow-timescale mode from nonlinear measurements, mirrors the trend of *τ*_1_, indicating that as the network becomes more contractile and reinforced, stress relaxation becomes progressively slower. As such, these results indicate a clear hierarchy of relaxation behaviors across linear and nonlinear regimes. The short-timescale modes–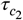 and *τ*_1_–are moderately sensitive to mesoscale architecture, with *τ*_1_ being more strongly affected by motor-induced changes. The intermediate mode *τ*_2_ reflects delayed relaxation in increasingly contractile networks, while the longest mode, 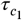, remains largely unaffected by structural reorganization. This separation of timescales indicates the presence of distinct mechanical regimes within actomyosin networks, wherein short-timescale modes capture local fluidization, and long-timescale modes preserve global mechanical memory.

To quantitatively connect network morphology to emergent mechanical properties, we analyzed the dependence of the plateau modulus *G*^0^ (black squares), zero-shear viscosity *η*^0^ (gray circles), and residual force *F*_*R*_ (gray downward triangles) on the structural correlation length *ξ*_*z*_ (Fig. 6B). All three mechanical parameters exhibit non-monotonic trends, first decreasing slightly and then increasing with *ξ*_*z*_, but with distinct sensitivities and scaling behaviors. The plateau modulus *G*^0^, which reflects the network’s elastic stiffness in the linear regime, initially softens as *ξ*_*z*_ increases from 0.65 to 1.0 *μ*m, but then rises steadily at larger *ξ*_*z*_, consistent with enhanced filament bundling or load-bearing connectivity in more coarsened architectures. In contrast, both *η*^0^ and *F*_*R*_, which reflect dissipative and stress-retentive properties, respectively, exhibit a more pronounced increase beyond *ξ*_*z*_ ∼ 1.2 *μ*m, following an initial dip. This sharp rise suggests that larger-scale contractile remodeling enhances viscous resistance and promotes the buildup of persistent internal stresses. While *G*^0^, *η*^0^, and *F*_*R*_ quantify distinct mechanical behaviors, their shared non-monotonic trends with increasing *ξ*_*z*_ suggest that all three are governed by common underlying structural transitions– from a loosely connected mesh to a contractile, bundled architecture. This coupling allows both stiffness and dissipative capacity to rise concurrently in highly remodeled networks. These results reveal that actomyosin-driven structural coarsening differentially modulates viscoelastic parameters: network stiffening (via *G*^0^) emerges gradually, while nonlinear stress storage and dissipation (via *F*_*R*_ and *η*^0^) respond more strongly to motor-induced mesoscale restructuring. These changes reinforce the network’s ability to resist deformation and maintain mechanical memory over extended timescales.

To complement the analysis based on spatial correlation length *ξ*_*z*_, we examined how key mechanical parameters vary as a function of the temporal correlation length *ξ*_*t*_, extracted from time-lapse confocal image sequences (SI Fig. S3). The data reveal divergent trends among plateau modulus *G*^0^, residual force *F*_*R*_, and zero-shear viscosity *η*^0^, suggesting distinct sensitivities to the temporal organization of network structure. Specifically, the zero-shear viscosity *η*^0^ exhibits a prominent peak at the lowest *ξ*_*t*_ (∼ 0.22 *μ*m), followed by a marked decline at intermediate values, and then a partial recovery at the largest *ξ*_*t*_ (∼ 0.62 *μ*m). This non-monotonic behavior implies that stress dissipation is maximized when dynamic rearrangements occur over shorter spatial scales, consistent with a fluid-like, actively remodeling network. In contrast, the residual force *F*_*R*_, a marker of internal stress retention, rises gradually with *ξ*_*t*_, peaking near *ξ*_*t*_ ∼ 0.60 *μ*m. This suggests that networks with more persistent structural features are better able to maintain tension over time, reflecting contractile stabilization. The plateau modulus *G*^0^, which characterizes elastic stiffness in the linear regime, shows a shallow minimum at intermediate *ξ*_*t*_, with elevated values at both lower and higher ends of the range. This biphasic trend indicates that stiffness can emerge both from dynamic filament entanglement in fluid-like states and from mechanically reinforced domains in more stabilized networks.

Collectively, our structural and mechanical measurements reveal a biphasic transition modulated by motor concentration: initial network fluidization at low *R*_MA_ gives way to stiffening and stress retention at high *R*_MA_, underpinned by progressive architectural coarsening. We now turn to a broader discussion of the mechanistic implications of these findings.

## 4 DISCUSSION

Our results demonstrate that increasing myosin motor content fundamentally alters both the structure and mechanics of reconstituted actin networks, driving a transition from a compliant, fluid-like state to a stiff, contractile architecture. We interpret these findings through the lens of motor-induced structural remodeling, microrheological signatures of fluidization versus reinforcement, nonlinear strain stiffening, and persistent stress retention in active cytoskeletal materials.

To understand this mechanical transformation, we first examined the network architecture using confocal fluorescence imaging, which revealed dramatic reorganization with increasing *R*_MA_ (Fig. 2A–B). In the absence of myosin, actin forms a homogeneous, isotropic mesh. As motor concentration increases (*R*_MA_ = 0.01–0.02), the network exhibits subtle heterogeneities indicative of nascent contractile remodeling. At higher myosin levels (*R*_MA_ = 0.04 and 0.08), pronounced coarsening occurs, giving rise to thick bundles, aster-like foci, and voids—structures characteristic of contractile superstructures [20, 61].

This structural evolution is quantitatively captured through spatial correlation analysis (Fig. 2C–E), which reveals monotonic growth of the axial correlation length *ξ*_z_ and saturation of the transverse correlation length *ξ*_t_. These trends suggest that while large-scale domains continue to grow under increasing tension, lateral reorganization reaches a steady state, reflecting stabilization of mesoscale structural motifs.

Mechanical measurements using linear optical tweezers microrheology further reinforce the picture of motor-induced structural modulation. The plateau modulus *G*^0^ and zero-shear viscosity *η*^0^ exhibit a non-monotonic dependence on *R*_MA_ (Fig. 3D–E). Low myosin levels lead to fluidization, evidenced by decreases in both *G*^0^ and *η*^0^, consistent with motor-induced transient disruption of filament connectivity. However, further increases in motor content reverse this trend: both parameters rise significantly, reflecting reinforcement through filament bundling and enhanced network connectivity. These observations highlight how motor content can dynamically tune viscoelastic properties, allowing for cellular modulation of mechanical states [37, 15].

Complementary nonlinear microrheology data underscore this transition, revealing pronounced strain stiffening at elevated motor concentrations (Fig. 4). The increase in both terminal stiffness (*K*_t_) and peak differential modulus (*K*_max_) with *R*_MA_ reflects enhanced force resistance, consistent with greater prestress and filament connectivity. Notably, the time to reach peak stiffness 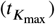 exhibits a non-monotonic dependence on motor content, suggesting that intermediate myosin levels optimize the kinetics of structural reinforcement.

Additionally, strain stiffening is followed by softening during extended deformation. This characteristic nonlinear mechanical response—initial stress stiffening followed by softening—has been widely reported in studies of cross-linked actin networks, where softening is typically attributed to force-induced unbinding of cross-linking proteins [52, 72, 71, 76, 77]. Similar behavior has been observed in biotin-streptavidin cross-linked microtubule networks [72], where initial stiffening gave way to plastic dissipation and sustained elasticity. Simulations from that study attributed the softening to crosslinker unbinding and network rearrangement and persistent elasticity to crosslinker rebinding. Actomyosin networks assembled with transient *α*-actinin crosslinkers are expected to behave similarly. Here, structural yielding may result from a combination of crosslinker unbinding and myosin-driven filament sliding, while rapid reattachment of *α*-actinin helps stabilize contractile structures as they form. The observed increase in *K*_max_ with higher *R*_MA_ further supports the idea that motor-induced prestress enhances the initial stiffness of the network prior to mechanical softening.

To further elucidate the mechanisms underlying the observed nonlinear mechanical responses, we examined force relaxation dynamics across motor concentrations in both the linear and nonlinear regimes (Fig. 6A). As such, we identified two dominant relaxation timescales: a fast component—denoted 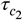 in the linear regime and *τ*_1_ in the nonlinear regime—and a slower component—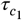 and *τ*_2_, respectively.

The fast relaxation mode aligns closely with theoretical predictions for actin entanglement dynamics. Specifically, the entanglement time *τ*_*e*_ for semiflexible filaments is expected to scale with the mesh size *ζ* and the persistence length *l*_*p*_ as 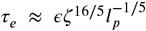, where *ϵ* is a friction coefficient estimated to be ∼3 s *μ*m^3^ for actin [53, 78]. Using *l*_*p*_ = 10 *μ*m for phalloidin-stabilized filaments, we estimate *τ*_*e*_ ≈ 0.33 s—comparable to our measured values of *τ*_1_ = 0.48 s and 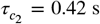. This agreement suggests that the fast relaxation mode reflects entanglement-scale dynamics. However, the elevated values of *τ*_1_ and 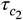 in passive networks indicate contributions beyond pure mesh confinement, such as filament rearrangements or *α*-actinin unbinding. As myosin is introduced, *τ*_1_ decreases toward *τ*_*e*_ and stabilizes at high *R*_MA_, consistent with enhanced local remodeling driven by motor-induced bundling and prestress. These results further suggest that fast relaxation becomes increasingly dominated by entanglement-mediated stress release in stiffened, actively remodeled networks.

In contrast, the slow relaxation timescales–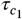 in the linear regime and *τ*_2_ in the nonlinear regime–exhibit weaker sensitivity to myosin concentration. In passive networks, *τ*_2_ ranges from 8–10 s and declines modestly to approximately 5–6 s with increasing *R*_MA_. This trend indicates that *τ*_2_ likely captures larger-scale processes such as filament sliding and global network reorganization. Meanwhile, 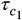 remains nearly constant across motor concentrations, suggesting that it reflects architecture-independent, stress-bearing modes that are robust to changes in active connectivity. The modest decrease in *τ*_2_ with myosin addition suggests that this mode is primarily governed by transient crosslinker turnover, rather than by motor-driven restructuring of the network. This interpretation is supported by prior findings that identify *α*-actinin unbinding kinetics as a key determinant of long-timescale stress relaxation in reconstituted actin networks [79].

Compared to other actin-based systems, the relaxation timescales measured here are generally faster than those observed in purely entangled or statically crosslinked networks [67, 66]. This distinction underscores the combined influence of transient crosslinking and active remodeling in accelerating relaxation dynamics. While *τ*_2_ corresponds well with modes previously associated with constraint release or global filament mobility, *τ*_1_ shows a pronounced dependence on motor activity and converges toward the theoretical entanglement time *τ*_*e*_ under conditions of high motor density. The results indicate a division of labor between two relaxation modes: *τ*_1_ represents local, motor-sensitive viscoelastic relaxation at the mesh scale, while *τ*_2_ reflects slower network-level remodeling influenced by crosslinker kinetics. This demonstrates how myosin motors adjust cytoskeletal mechanics by coordinating structural organization and temporal response across various physical regimes.

To elucidate how mesoscale structure modulates relaxation dynamics and viscoelastic function in actomyosin networks, we examined the dependence of relaxation timescales and mechanical parameters on both *ξ*_*z*_ and *ξ*_*t*_ correlation lengths (Fig. 6, SI Fig. S3). As shown in Fig. 6(A), both *τ*_1_ and *τ*_2_ exhibit non-monotonic dependencies on *ξ*_*z*_, with clear minima near *ξ*_*z*_ ∼ 1 *μ*m. These minima indicate a regime of enhanced dynamic remodeling and faster stress relaxation, suggesting reduced network cohesion or partial disruption of load-bearing structures at this intermediate scale. In contrast, 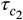 exhibits a slight increase with *ξ*_*z*_ until it reaches maximum at intermediate *ξ*_*z*_, pointing to a transient slowdown in co-operative or collective rearrangements. The opposing trends of *τ*_1_, *τ*_2_ versus 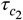 highlight the coexistence of accelerated local relaxation and temporarily arrested collective modes in partially coarsened networks. Meanwhile, 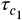 remains relatively invariant across *ξ*_*z*_, suggesting it reflects long-timescale processes largely unaffected by structural correlation, such as crosslinker unbinding or large-scale network reorganization.

Analysis of the mechanical response reveals that the plateau modulus *G*^0^ and zero-shear viscosity *η*^0^ vary non-monotonically with the longitudinal correlation length *ξ*_*z*_, each reaching a clear minimum at *ξ*_*z*_ ∼ 1 *μ*m. In contrast, the residual force *F*_*R*_ increases gradually across the entire range of *ξ*_*z*_. This regime of mechanical softening coincides with the minima observed in the fast relaxation timescales *τ*_1_ and *τ*_2_, collectively indicating that networks with intermediate *ξ*_*z*_ are more compliant and dynamically fluid. At larger *ξ*_*z*_, all three mechanical parameters–*G*^0^, *η*^0^, and *F*_*R*_–increase, signaling a transition to contractile, bundled architectures that resist deformation and maintain long-lived internal stress. The concurrent rise in *η*^0^ and *F*_*R*_ at high *ξ*_*z*_ underscores enhanced energy dissipation and stress memory associated with motor-driven structural reinforcement.

A similar structure–function coupling is observed with transverse correlation length *ξ*_*t*_ (SI Fig. S3). Although the trends are less pronounced, both *τ*_1_ and 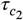 again display non-monotonic behaviors, consistent with a dynamically heterogeneous regime at intermediate *ξ*_*t*_. Notably, 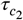 is particularly sensitive to transverse structural changes. Meanwhile, mechanical parameters *G*^0^ and *F*_*R*_ increase monotonically with *ξ*_*t*_, while *η*^0^ peaks at intermediate values—again supporting the interpretation that viscoelastic dissipation is maximized during partial remodeling, whereas fully coarsened networks exhibit enhanced mechanical reinforcement.

## 5 CONCLUSION

The ability of cytoskeletal networks to dynamically modulate their mechanical properties is fundamental to a wide range of cellular processes. Our study demonstrates that myosin motor activity acts as a potent regulator of both structural organization and mechanical behavior in reconstituted actin networks. By systematically varying the motor-to-actin molar ratio (*R*_MA_ = 0–0.08) at a fixed concentration of the transient crosslinker *α*-actinin (0.05), we reveal a dynamic transition from fluid-like to contractile, stress-bearing architectures. This transition is characterized by non-monotonic changes in both viscoelastic properties and network morphology, including an initial decrease followed by an increase in plateau modulus (*G*^0^) and viscosity (*η*^0^), a minimum in fast relaxation time (*τ*_1_) near intermediate spatial correlation lengths, and a peak in strain-induced stiffening timescale 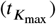, all reflecting the competing effects of motor-driven fluidization and contractile stabilization. Our results further identify two mechanistically distinct relaxation modes–*τ*_1_ and *τ*_2_–that remain consistently separated across motor conditions, indicating fundamentally different underlying mechanisms. While both relaxation times show only modest variation with increasing *R*_MA_, particularly at low motor concentrations, their persistence as distinct timescales highlights the coexistence of multiple stress dissipation pathways. The fast mode *τ*_1_ likely reflects local rearrangements at the mesh scale, influenced by filament connectivity and bundling, whereas the slower mode *τ*_2_ is governed by the unbinding and rebinding dynamics of the transient *α*-actinin crosslinks resulting in large-scale structural relaxation. This separation of timescales highlights the multiscale nature of stress regulation in cytoskeletal networks, revealing how passive crosslinking and active motor activity operate in parallel to shape the mechanical response. By coupling structure and mechanics across scales, this study provides a quantitative framework for understanding how cells modulate cytoskeletal behavior through motor activity.

## Authors contributions

BI and MT performed experiments, analyzed data, and drafted the manuscript; BJG designed and guided experiments, analyzed and interpreted data, and wrote the manuscript.

## Acknowledgements

We thank Bucknell University for the financial support.

## Data Availability Statement

The data presented in this study are available on request from the corresponding author.

## References

[1] Herrmann H, Bar H, Kreplak L, Strelkov SV, Aebi U. Intermediate filaments: from cell architecture to nanomechanics. Nature Reviews Molecular Cell Biology 2007; 8(7): 562–573. http://dx.doi.org/10.1038/nrm2197 doi: 10.1038/nrm2197

[2] Alberts B, Johnson A, Lewis J, Raff M, Roberts K, Walter P. Molecular biology of the cell. 6. Garland Science. 6th ed. 2017.

[3] Pollard TD, Borisy GG. Cellular motility driven by assembly and disassembly of actin filaments. Cell 2003; 112(4): 453–465. http://dx.doi.org/10.1016/S0092-8674(03)00076-2 doi: 10.1016/S0092-8674(03)00076-2

[4] Bray D. Cell movements: from molecules to motility. Garland Science. 2001.

[5] Ridley AJ, Schwartz MA, Burridge K, et al. Cell migration: integrating signals from front to back. Science 2003; 302(5651): 1704–1709.

[6] Papakonstanti E, Stournaras C. Cell responses regulated by early reorganization of actin cytoskeleton. FEBS letters 2008; 582(14): 2120–2127.

[7] Heng YW, Koh CG. Actin cytoskeleton dynamics and the cell division cycle. The international journal of biochemistry & cell biology 2010; 42(10): 1622–1633.

[8] Fletcher DA, Mullins RD. Cell mechanics and the cytoskeleton. Nature 2010; 463(7280): 485–492. http://dx.doi.org/10.1038/nature08908 doi: 10.1038/nature08908

[9] Blaser H, Reichman-Fried M, Castanon I, et al. Migration of zebrafish primordial germ cells: a role for myosin contraction and cytoplasmic flow. Developmental Cell 2006; 11(5): 613–627. http://dx.doi.org/10.1016/j.devcel.2006.09.023 doi: 10.1016/j.devcel.2006.09.023

[10] Paluch EK, Aspalter IM, Sixt M. Focal adhesion– independent cell migration. Annual review of cell and developmental biology 2016; 32(1): 469–490.

[11] Lieleg O, Schmoller KM, Cyron CJ, Luan Y, Wall WA, Bausch AR. Structural polymorphism in heterogeneous cytoskeletal networks. Soft Matter 2009; 5(9): 1796–1803.

[12] Blanchoin L, Boujemaa-Paterski R, Sykes C, Plastino J. Actin dynamics, architecture, and mechanics in cell motility. Physiological reviews 2014; 94(1): 235–263.

[13] Mattila PK, Lappalainen P. Filopodia: molecular architecture and cellular functions. Nature Reviews Molecular Cell Biology 2008; 9(6): 446–454. http://dx.doi.org/10.1038/nrm2406 doi: 10.1038/nrm2406

[14] Vakhrusheva A, Murashko A, Trifonova E, Efremov YM, Timashev P, Sokolova O. Role of actin-binding proteins in the regulation of cellular mechanics. European Journal of Cell Biology 2022; 101(3): 151241.

[15] Murrell MP, Gardel ML. F-actin buckling coordinates contractility and severing in a biomimetic actomyosin cortex. Proceedings of the National Academy of Sciences 2012; 109(51): 20820–20825.

[16] Gurmessa B, Francis M, Rust MJ, Das M, Ross JL, Robertson-Anderson RM. Counterion crossbridges enable robust multiscale elasticity in actin networks. Physical Review Research 2019; 1(1): 013016.

[17] Deshpande S, Pfohl T. Real-time dynamics of emerging actin networks in cell-mimicking compartments. PloS one 2015; 10(3): e0116521.

[18] Tang JX, Janmey PA. The Polyelectrolyte Nature of Factin and the Mechanism of Actin Bundle Formation (). Journal of Biological Chemistry 1996; 271(15): 8556–8563.

[19] Huber F, Strehle D, Käs J. Counterion-induced formation of regular actin bundle networks. Soft Matter 2012; 8(4): 931–936.

[20] Murrell M, Oakes PW, Lenz M, Gardel ML. Forcing cells into shape: the mechanics of actomyosin contractility. Nature reviews Molecular cell biology 2015; 16(8): 486–498.

[21] Mizuno D, Tardin C, Schmidt CF, MacKintosh FC. Nonequilibrium mechanics of active cytoskeletal networks. Science 2007; 315(5810): 370–373.

[22] Koenderink G, Atakhorrami M, MacKintosh F, Schmidt CF. High-frequency stress relaxation in semiflexible polymer solutions and networks. Physical review letters 2006; 96(13): 138307.

[23] Gardel M, Nakamura F, Hartwig J, Crocker JC, Stossel T, Weitz D. Stress-dependent elasticity of composite actin networks as a model for cell behavior. Physical review letters 2006; 96(8): 088102.

[24] Mukhina S, Wang Yl, Murata-Hori M. ??-Actinin is required for tightly regulated remodeling of the actin cortical network during cytokinesis. Developmental cell 2007; 13(4): 554–565.

[25] Hotulainen P, Lappalainen P. Stress fibers are generated by two distinct actin assembly mechanisms in motile cells. The Journal of Cell Biology 2006; 173(3): 383–394.

[26] Dwyer ME, Robertson-Anderson RM, Gurmessa BJ. Nonlinear microscale mechanics of actin networks governed by coupling of filament crosslinking and stabilization. Polymers 2022; 14(22): 4980.

[27] Courson DS, Rock RS. Actin cross-link assembly and disassembly mechanics for ??-actinin and fascin. Journal of Biological Chemistry 2010; 285(34): 26350–26357.

[28] Lee G, Leech G, Rust MJ, et al. Myosin-driven actin-microtubule networks exhibit self-organized contractile dynamics. Science Advances 2021; 7(6): eabe4334.

[29] Gardel ML, Nakamura F, Hartwig JH, Crocker JC, Stossel TP, Weitz DA. Prestressed F-actin networks cross-linked by hinged filamins replicate mechanical properties of cells. Proceedings of the National Academy of Sciences 2006; 103(6): 1762–1767.

[30] Seetharaman S, Etienne-Manneville S. Cytoskeletal crosstalk in cell migration. Trends in cell biology 2020; 30(9): 720–735.

[31] Ideses Y, Sonn-Segev A, Roichman Y, Bernheim-Groswasser A. Myosin II does it all: assembly, remodeling, and disassembly of actin networks are governed by myosin II activity. Soft Matter 2013; 9(29): 7127–7137.

[32] Betapudi V. Life without double-headed non-muscle myosin II motor proteins. Frontiers in chemistry 2014; 2: 45.

[33] Silva S. eM, Depken M, Stuhrmann B, Korsten M, MacKintosh FC, Koenderink GH. Active multistage coarsening of actin networks driven by myosin motors. Proceedings of the National Academy of Sciences 2011; 108(23): 9408–9413.

[34] Laevsky G, Knecht DA. Cross-linking of actin filaments by myosin II is a major contributor to cortical integrity and cell motility in restrictive environments. Journal of Cell Science 2003; 116(18): 3761–3770.

[35] Wachsstock DH, Schwarz W, Pollard T. Cross-linker dynamics determine the mechanical properties of actin gels. Biophysical journal 1994; 66(3): 801–809.

[36] Humphrey D, Duggan C, Saha D, Smith D, Käs J. Active fluidization of polymer networks through molecular motors. Nature 2002; 416(6879): 413–416.

[37] Koenderink GH, Dogic Z, Nakamura F, et al. An active biopolymer network controlled by molecular motors. Proceedings of the National Academy of Sciences 2009; 106(36): 15192–15197.

[38] Reichl EM, Ren Y, Morphew MK, et al. Interactions between myosin and actin crosslinkers control cytokinesis contractility dynamics and mechanics. Current Biology 2008; 18(7): 471–480.

[39] Martens JC, Radmacher M. Softening of the actin cytoskeleton by inhibition of myosin II. Pflügers Archiv-European Journal of Physiology 2008; 456(1): 95–100.

[40] Rauzi M, Verant P, Lecuit T, Lenne PF. Nature and anisotropy of cortical forces orienting Drosophila tissue morphogenesis. Nature Cell Biology 2008; 10(12): 1401–1410.

[41] Huxley H. Fifty years of muscle and the sliding filament hypothesis. European Journal of Biochemistry 2004; 271(8): 1403–1415.

[42] Thoresen T, Lenz M, Gardel ML. Reconstitution of contractile actomyosin bundles. Biophysical Journal 2011; 100(11): 2698–2705.

[43] Matsuda K, Jung W, Sato Y, et al. Myosin-induced Factin fragmentation facilitates contraction of actin networks. Cytoskeleton 2024; 81(8): 339–355.

[44] Lieleg O, Claessens MM, Heussinger C, Frey E, Bausch AR. Mechanics of bundled semiflexible polymer networks. Physical review letters 2007; 99(8): 088102.

[45] Gardel ML, Shin JH, MacKintosh FC, Mahadevan L, Matsudaira P, Weitz DA. Elastic behavior of cross-linked and bundled actin networks. Science 2004; 304(5675): 1301–1305.

[46] Broedersz CP, MacKintosh FC. Modeling semiflexible polymer networks. Reviews of Modern Physics 2014; 86(3): 995.

[47] Storm C, Pastore JJ, MacKintosh FC, Lubensky TC, Janmey PA. Nonlinear elasticity in biological gels. Nature 2005; 435(7039): 191–194.

[48] Broedersz C, MacKintosh F. Molecular motors stiffen non-affine semiflexible polymer networks. Soft Matter 2011; 7(7): 3186–3191.

[49] Kasza K, Broedersz C, Koenderink G, et al. Actin filament length tunes elasticity of flexibly cross-linked actin networks. Biophysical journal 2010; 99(4): 1091–1100.

[50] Schmoller K, Fernandez P, Arevalo R, Blair D, Bausch A. Cyclic hardening in bundled actin networks. Nature communications 2010; 1(1): 134.

[51] Bendix PM, Koenderink GH, Cuvelier D, et al. Actin tracks and trails from a diffusing particle in a branched actin network. Biophysical Journal 2008; 94(8): 3126–3136.

[52] Tharmann R, Claessens MM, Bausch AR. Viscoelasticity of isotropically cross-linked actin networks. Physical review letters 2007; 98(8): 088103.

[53] Isambert H, Maggs A. Dynamics and rheology of actin solutions. Macromolecules 1996; 29(3): 1036–1040.

[54] Gardel M, Valentine M, Crocker JC, Bausch A, Weitz D. Microrheology of entangled F-actin solutions. Physical review letters 2003; 91(15): 158302.

[55] Schmidt CF, Baermann M, Isenberg G, Sackmann E. Chain dynamics, mesh size, and diffusive transport in networks of polymerized actin: a quasielastic light scattering and microfluorescence study. Macromolecules 1989; 22(9): 3638–3649.

[56] Weigand W, Messmore A, Tu J, et al. Active microrheology determines scale-dependent material properties of Chaetopterus mucus. PloS one 2017; 12(5): e0176732.

[57] Chapman CD, Lee K, Henze D, Smith DE, Robertson-Anderson RM. Onset of non-continuum effects in microrheology of entangled polymer solutions. Macro-molecules 2014; 47(3): 1181–1186.

[58] Francis ML, Ricketts SN, Farhadi L, et al. Non-monotonic dependence of stiffness on actin crosslinking in cytoskeleton composites. Soft Matter 2019; 15(44): 9056–9065.

[59] Gurmessa BJ, Rust MJ, Das M, Ross JL, Robertson-Anderson RM. Salt-mediated stiffening, destruction, and resculpting of actomyosin network. Frontiers in Physics 2021: 667.

[60] Robertson C, George SC. Theory and practical recommendations for autocorrelation-based image correlation spectroscopy. Journal of biomedical optics 2012; 17(8): 080801.

[61] Sheung JY, Achiriloaie DH, Currie C, et al. Motor-Driven Restructuring of Cytoskeleton Composites Leads to Tunable Time-Varying Elasticity. ACS Macro Letters 2021; 10: 1151–1158.

[62] Brau R, Ferrer J, Lee H, et al. Passive and active microrheology with optical tweezers. Journal of Optics A: Pure and Applied Optics 2007; 9(8): S103.

[63] Robertson-Anderson RM. Optical tweezers microrheology: from the basics to advanced techniques and applications. 2018.

[64] Tassieri M, Evans R, Warren RL, Bailey NJ, Cooper JM. Microrheology with optical tweezers: data analysis. New Journal of Physics 2012; 14(11): 115032.

[65] Falzone TT, Blair S, Robertson-Anderson RM. Entangled F-actin displays a unique crossover to microscale nonlinearity dominated by entanglement segment dynamics. Soft matter 2015; 11(22): 4418–4423.

[66] Gurmessa B, Ricketts S, Robertson-Anderson RM. Nonlinear Actin Deformations Lead to Network Stiffening, Yielding, and Nonuniform Stress Propagation. Biophysical Journal 2017; 113(7): 1540–1550.

[67] Gurmessa B, Fitzpatrick R, Falzone TT, Robertson-Anderson RM. Entanglement Density Tunes Microscale Nonlinear Response of Entangled Actin. Macromolecules 2016.

[68] Bendix PM, Koenderink GH, Cuvelier D, et al. A quantitative analysis of contractility in active cytoskeletal protein networks. Biophysical journal 2008; 94(8): 3126–3136.

[69] Luan Y, Lieleg O, Wagner B, Bausch AR. Micro- and macrorheological properties of isotropically cross-linked actin networks. Biophysical journal 2008; 94(2): 688–693.

[70] Schmidt FG, Hinner B, Sackmann E. Microrheometry underestimates the values of the viscoelastic moduli in measurements on F-actin solutions compared to macrorheometry. Physical Review E 2000; 61(5): 5646.

[71] Kim T, Gardel ML, Munro E. Determinants of fluidlike behavior and effective viscosity in cross-linked actin networks. Biophysical journal 2014; 106(3): 526–534.

[72] Yang Y, Bai M, Klug WS, Levine AJ, Valentine MT. Microrheology of highly crosslinked microtubule networks is dominated by force-induced crosslinker unbinding. Soft matter 2013; 9(2): 383–393.

[73] Lieleg O, Schmoller K, Claessens MMAE, Bausch AR. Cytoskeletal polymer networks: viscoelastic properties are determined by the microscopic interaction potential of cross-links. Biophysical journal 2009; 96(11): 4725–4732.

[74] Doi M, Edwards SF. The theory of polymer dynamics. 73. oxford university press. 1988.

[75] De Gennes PG. Scaling concepts in polymer physics. Cornell university press. 1979.

[76] Abhilash A, Purohit PK, Joshi SP. Stochastic rate-dependent elasticity and failure of soft fibrous networks. Soft Matter 2012; 8(26): 7004–7016.

[77] Plagge J, Fischer A, Heussinger C. Viscoelasticity of reversibly crosslinked networks of semiflexible polymers. Physical Review E 2016; 93(6): 062502.

[78] Käs J, Strey H, Tang J, et al. F-actin, a model polymer for semiflexible chains in dilute, semidilute, and liquid crystalline solutions.. Biophysical journal 1996; 70(2): 609.

[79] Lieleg O, Claessens MM, Bausch AR. Slow dynamics and internal stress relaxation in bundled cytoskeletal networks. Physical Review Letters 2008; 101(10): 108101.

