## Supplemental figures for "Motor-driven modulation of actin network mechanics across linear and nonlinear regimes"

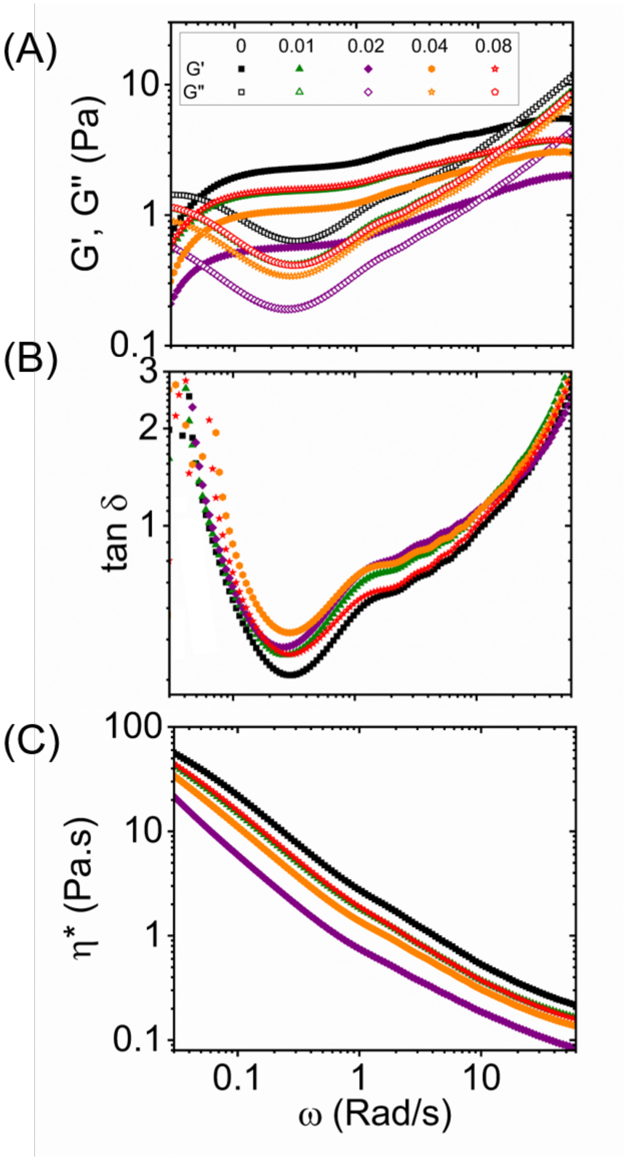


SI Fig. S1: (A) Storage modulus $G'$ (solid markers) and loss modulus $G''$ (open markers) plotted as a function of angular frequency $\omega$ for actomyosin networks with increasing myosin-to-actin molar ratio $R_{MA}$. The data demonstrate that low levels of myosin reduce both $G'$ and $G''$, consistent with network fluidization, while higher motor concentrations recover and enhance elasticity. (B) Loss tangent $\tan\delta = G''/G'$ versus frequency highlights a non-monotonic dependence on $R_{MA}$, with minimum $\tan\delta$ shifting across motor concentrations, reflecting changes in network dissipation. (C) Complex viscosity $\eta^*(\omega) = \sqrt{G'^2 + G''^2}/\omega$ decreases with frequency for all conditions, as expected for viscoelastic systems, and exhibits a minimum near intermediate motor levels, consistent with ATP-driven softening followed by mechanical reinforcement.

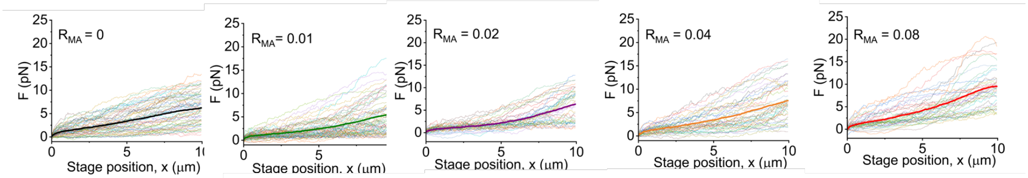


SI Fig. S2: Individual raw force–displacement traces ($F(x)$) obtained from optical tweezers experiments across five myosin-to-actin molar ratios ($R_{MA}$ = 0, 0.01, 0.02, 0.04, 0.08). Each curve represents a single trial with a different microsphere probe in an independent region of the sample chamber. Force responses were measured as the stage displaced the bead through the actin-myosin network at a constant speed of 5 $\mu$m/s. Each panel shows all trials corresponding to a specific $R_{MA}$ condition, with two replicates of 25 trials each and displayed together. These data underpin the ensemble-averaged curves shown in Fig. 5A of the main text.

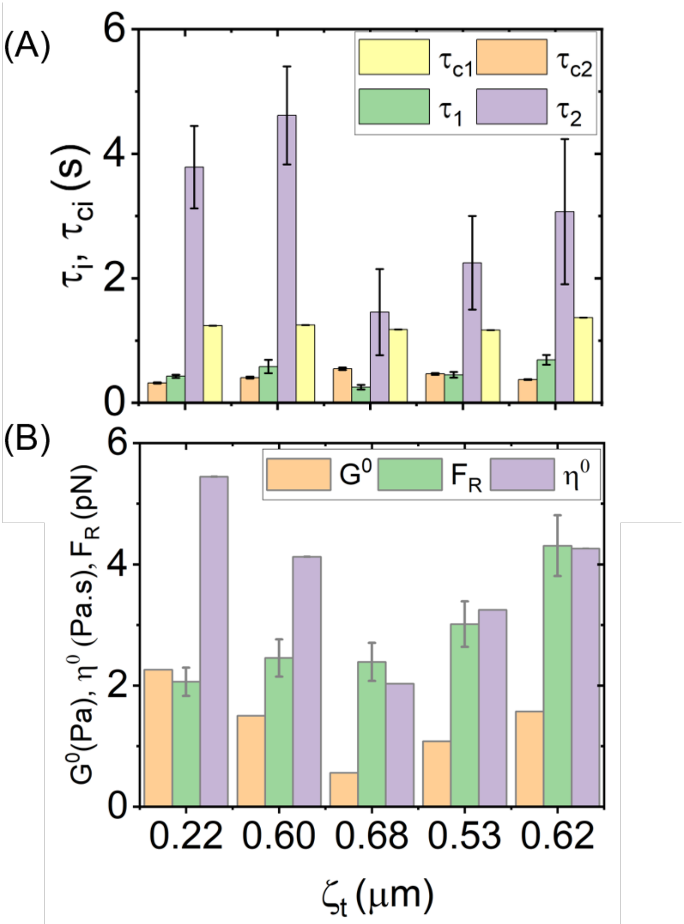


**SI Fig. S3: Mechanical properties of actomyosin networks as a function of temporal correlation length $\xi_t$.** (A) Relaxation time constants extracted from linear and nonlinear microrheology plotted against $\xi_t$, the temporal structural correlation length obtained from time-lapse confocal projections. Shown are fast and slow relaxation times $\tau_1$ (green) and $\tau_2$ (purple) from nonlinear microrheology, and $\tau_{c_1}$ (yellow) and $\tau_{c_2}$ (orange) from linear microrheology. $\tau_1$ remains relatively stable across $\xi_t$, while $\tau_2$ exhibits a strong peak at low $\xi_t$, indicating enhanced dissipation in more dynamically heterogeneous networks. The slowest linear timescale $\tau_{c_1}$ remains largely invariant, suggesting that long-lived stress-bearing modes are insensitive to temporal reorganization. In contrast, $\tau_{c_2}$ displays moderate variation, peaking at intermediate $\xi_t$. (B) Key viscoelastic and mechanical parameters as a function of $\xi_t$. The plateau modulus $G^0$ (orange) and residual force $F_R$ (green) show shallow minima at intermediate $\xi_t$, while zero-shear viscosity $\eta^0$ (purple) exhibits a pronounced peak at $\xi_t \sim 0.22$ $\mu$m, followed by a significant decline. These trends suggest that networks with short-range dynamic rearrangements dissipate stress more efficiently, whereas those with persistent structures retain internal stress and display increased elasticity. Error bars represent the standard error across measurements.
